# The transcription factor Odd-paired regulates temporal identity in transit-amplifying neural progenitors via an incoherent feed-forward loop

**DOI:** 10.1101/576736

**Authors:** Merve Deniz Abdusselamoglu, Elif Eroglu, Thomas R. Burkard, Juergen A. Knoblich

## Abstract

Neural progenitors undergo temporal patterning to generate diverse neurons in a chronological order. This process is well-studied in the developing *Drosophila* brain and conserved in mammals. During larval stages, intermediate neural progenitors (INPs) serially express Dichaete (D), grainyhead (Grh) and eyeless (Ey/Pax6), but how the transitions are regulated is not precisely understood. Here, we developed a method to isolate transcriptomes of INPs in their distinct temporal states to identify a complete set of temporal patterning factors. Our analysis identifies odd-paired (opa), as a key regulator of temporal patterning. Temporal patterning is initiated when the SWI/SNF complex component Osa induces D and its repressor Opa at the same time but with distinct kinetics. Then, high Opa levels repress D to allow Grh transcription and progress to the next temporal state. We propose that Osa and its target genes opa and D form an incoherent feedforward loop (FFL) and a new mechanism allowing the successive expression of temporal identities.

## Introduction

During brain development, neural stem cells (NSCs) generate large numbers of highly diverse neuronal and glial cells in chronological order (Cepko, Austin, Yang, Alexiades, & Ezzeddine, 1996; Gao et al., 2014; Greig, Woodworth, Galazo, Padmanabhan, & Macklis, 2013; Holguera & Desplan, 2018). Through a phenomenon known as temporal patterning, NSCs acquire properties that change the fate of their progeny over time (Kohwi, Lupton, Lai, Miller, & Doe, 2013; Mattar, Ericson, Blackshaw, & Cayouette, 2015a; Okamoto et al., 2016). Importantly, temporal patterning of NSCs is an evolutionary conserved process and has been observed across species ranging from insects to mammals (Alsiö, Tarchini, Cayouette, & Livesey, 2013; Livesey & Cepko, 2001; Toma, Kumamoto, & Hanashima, 2014). During mammalian brain development, neural progenitors in the central nervous system (CNS) undergo temporal patterning by relying on both extrinsic as well as progenitor-intrinsic cues. Wnt7, for example, is an extracellular ligand required for the switch from early to late neurogenesis in cortical progenitors (W. Wang et al., 2016), Ikaros (the ortholog of the Drosophila Hunchback), in contrast, is an intrinsic factor specifying early-born neuronal fates (Mattar, Ericson, Blackshaw, & Cayouette, 2015b). Like Ikaros, intrinsic temporal identity factors in vertebrates are often homologous to factors described in *Drosophila* (Naka, Nakamura, Shimazaki, & Okano, 2008; Ren et al., 2017; Syed, Mark, & Doe, 2017). How these factors are involved in neuronal fate specification and how they are regulated remain unknown.

*Drosophila* has been crucial to understanding stem cell biological mechanisms and in particular distinct temporal patterning processes (Homem & Knoblich, 2012). During embryonic neurogenesis, *Drosophila* NSCs, called Neuroblasts (NBs), undergo temporal patterning through a cascade of transcription factors (Isshiki, Pearson, Holbrook, & Doe, 2001). During larval neurogenesis, NB temporal patterning relies on opposing gradients of two RNA-binding proteins (Liu et al., 2015; Syed et al., 2017). Temporal patterning is also seen in intermediate neural progenitors (INPs), the transit-amplifying progeny of a discrete subset of larval NBs called type II NBs (Bayraktar & Doe, 2013). Once they arise from an asymmetric division of a type II NB, newborn INPs undergo several maturation steps before they resume proliferation: they first turn on earmuff (erm) and Asense (ase) and finally Deadpan (Dpn) expression to become mature INPs (mINP) (Bello, Izergina, Caussinus, & Reichert, 2008; Boone & Doe, 2008; Bowman et al., 2008; Janssens et al., 2014; Walsh & Doe, 2017). mINPs again divide asymmetrically for 3-6 times to generate ganglion mother cells (GMCs), which in turn generate a pair of neurons or glia through a symmetric division. In analogy to embryonic NBs, temporal patterning in INPs was suggested to be mediated by a cascade of transcription factors (Isshiki et al., 2001). Indeed, the sequential expression of Dichaete (D), Grainyhead (Grh) and Eyeless (Ey) is required to generate different neurons: D+ INPs produce Brain-specific homeobox (Bsh)+ neurons, while Ey+ INPs produce Toy+ neurons (Bayraktar & Doe, 2013).

The three temporal identity factors are regulated through various regulatory interactions (Bayraktar & Doe, 2013; Doe, 2017): D is necessary, but not sufficient, for activating Grh. Grh instead is required for repression of D and activation of Ey (Bayraktar & Doe, 2013). Therefore, INP temporal patterning is thought to be regulated by a ‘feedforward activation and feedback repression’ mechanism (Fig1A). Intriguingly however, INP temporal patterning also critically requires the SWI/SNF chromatin remodeling complex subunit Osa (Eroglu et al., 2014). Although Osa is not considered a specific temporal identity factor, it is required to initiate temporal patterning by activating the initial factor D. While the Osa target gene hamlet is required for the Grh-to-Ey transition (Eroglu et al., 2014), regulation of the first transition is less well understood. This result suggests that in addition to feedforward activation and feedback repression, temporal switch genes are required to ensure correct INP temporal patterning. Nevertheless, D and ham double knock down (k.d.) phenotypes do not recapitulate the complete loss of temporal patterning initiation observed in Osa-depleted type II NB lineages, suggesting the contribution of additional unidentified factors.

**Figure 1.**
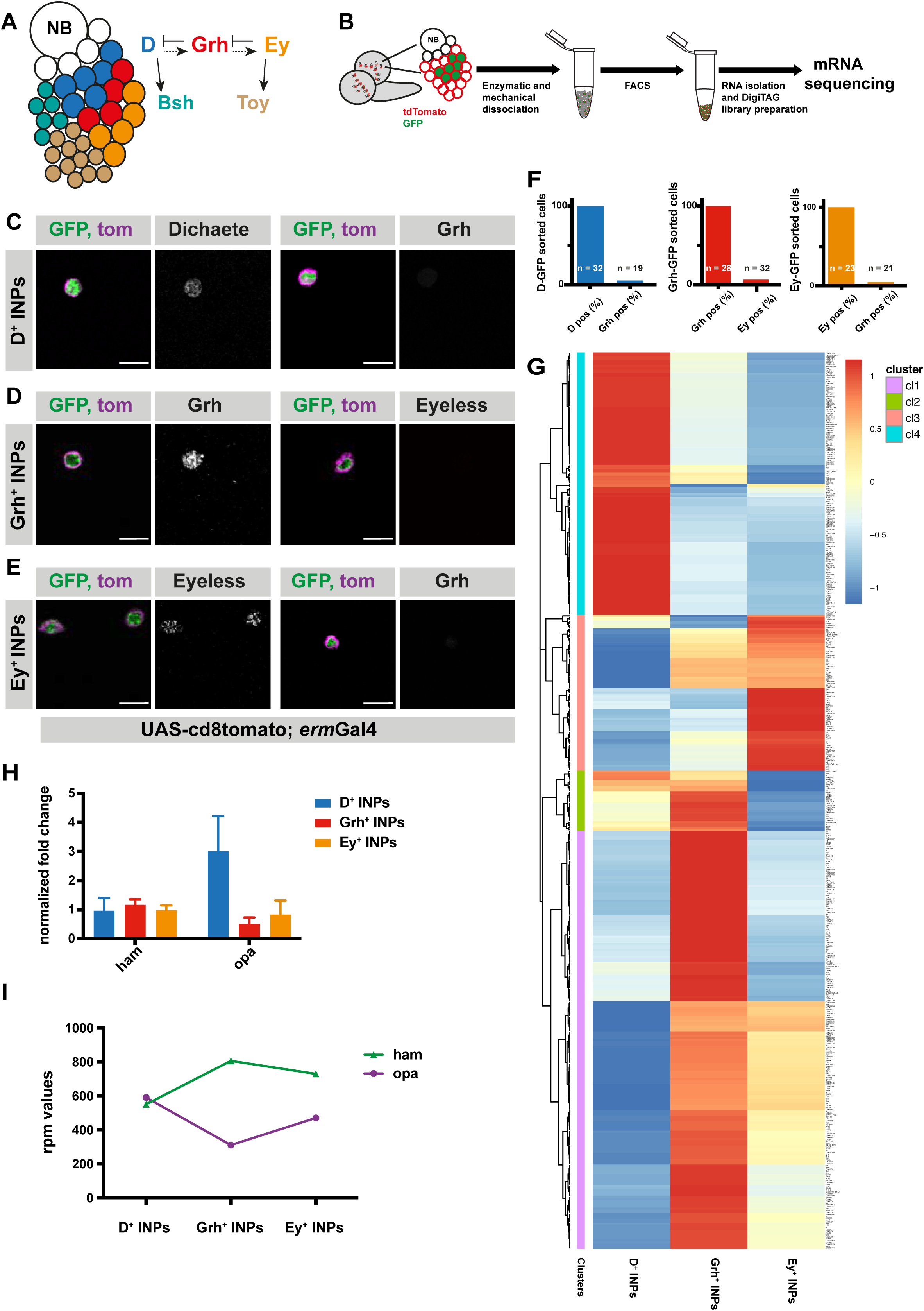
Transcriptomic analysis of temporally staged-INPs. (**A**) Cartoon depicting a typical type II neuroblast of larval *Drosophila* brain; NB and imINPs (empty circles) are followed by mINPs and neurons, GMCs omitted for simplicity. INPs are temporally patterned with Dichaete (blue), Grainyhead (red), and Eyeless (orange), and neurons are Bsh (green) or Toy (brown) positive. Summary of the regulation of temporal identity factors, and their progeny. **(B)** Cartoon illustrating the strategy used to isolate temporally-staged INPs. (**C-E**) D-, Grh and Ey-GFP FACS-sorted cells are stained for D and Grh (**C**), Grh or Ey (**D-E**), GFP-tagging temporal identity factors (in green, D or, Grh or Ey), tdTomato tagging the membrane of INPs (magenta), antibody staining (gray) scale bar 10 μm, (induced with ermGal4, marked with membrane bound tdTomato). (**F**) Graphs showing the percentage of temporal identity positive cells in D-, Grh-or Ey-GFP FACS sorted cells. n numbers are depicted on the graphs. (**G**) Hierarchical clustering analysis of gene log2fc between three different temporally-staged INP populations. **(H)** qPCR analysis of opa and ham expression levels in FACS-sorted D^+^, Grh^+^ and Ey^+^ INPs. Data are mean ± SD, n=3, genes were normalized to Act5c, and then the average expression levels, Delta-Delta Ct method is used. (**I**) Graph showing the rpm levels of opa and ham between different INP temporal stages.

Here, we describe a FACS-based method to isolate INPs from three different temporal identities. By comparing the transcriptomic profiles of each set of INPs, we identify odd-paired (opa), a direct target of Osa, as a regulator of temporal patterning and repressor of D. Though Osa enables both D and Opa expression, Opa’s slower activation kinetics allow D to function in a short time window before being repressed by Opa. This mode of action fits with an incoherent feedforward-loop (FFL) motif and uncovers a novel mechanism controlling temporal patterning during neurogenesis.

## Results

### Transcriptome analysis of distinct INP temporal states

To obtain a comprehensive list of temporally regulated genes in INPs, we used FACS-to purify INPs at each of their three temporal states: D^+^, Grh^+^ and Ey^+^ (Fig1B). For this, we generated fly lines expressing tdTomato under an INP specific promoter (erm-Gal4 > CD8::tdTomato) and expressing GFP-fusions of one of the temporal identity factors (D-GFP, Grh-GFP and Ey-GFP, FigS1A). After dissection and dissociation of third instar larval brains, GFP-positive INP populations (D-GFP^+^, Grh-GFP^+^ and Ey-GFP^+^) were identified (Fig1B and supp1B) as the largest cells with highest GFP and tdTomato expression (Fig1supp1B). Using immunofluorescence (IF), these cells were verified to be mature INPs (Fig1supp1C-D). All sorted cells within the INP populations expressed Dpn, indicating a 100% mature INP identity, while unsorted cells showed a mixture of Dpn^+^ and Dpn^-^ cells (Fig1supp1C-D). We validated the temporal identity of the progenitors by performing IF for their respective temporal identity markers (Fig1C-F and supp2A-C). Importantly, each GFP^+^ sorted INP population was 100% positive for its respective temporal marker (Fig1F). In contrast, the unsorted cells consisted of mixed cell populations containing various temporal identities (Fig1supp2B). Lastly, we tested for the presence of sorted cells expressing markers of two temporal identities, which reflects transition states of INP temporal patterning as occurs in vivo. Analyzing Grh IF on D-GFP^+^ and Ey-GFP^+^ sorted cells, and Ey IF on Grh-GFP^+^ sorted cells revealed that sorted populations contained only 4-6% of such double-positive cells (Fig1C-F, and Fig1supp2A-C), suggesting we can isolate almost pure populations of different temporal states. Collectively, we established the genetic tools and methodology to precisely sort INPs into separate populations according to their three distinct temporal states.

Since our stringent FACS sorting conditions led to low RNA yields, we generated cDNA libraries using DigiTag (Landskron et al., 2018; Wissel et al., 2018). With this RNA sequencing strategy, we found 458 genes expressed differently between D^+^ and Grh^+^ INPs, and 466 genes between Grh^+^ and Ey^+^ INPs (FDR 0.05, log2foldchange>1, and Rpm (reads per million mapped reads) > 10 in one of three samples/D^+^, Grh^+^ or Ey^+^ INPs). Hierarchical clustering identified genes specifically expressed in certain temporal states, and therefore potentially involved in temporal patterning (Fig1G). First, we confirmed the quality of our dataset by examining the transcriptional changes of temporal identity genes with quantitative PCR (qPCR) (Fig1supp1E). As expected, each temporal state had high expression levels of their own temporal identity genes. Second, we confirmed the expression of known temporal identity genes (Fig1supp1F). Third, we performed GO-term analysis on the identified gene clusters. Genes upregulated in D^+^ INPs showed enrichment for mitochondrial translation, cellular nitrogen compound metabolic process and gene expression (Fig1supp2F). Genes upregulated in Ey+ INPs were enriched for neurogenesis and sequence-specific DNA binding (Fig1supp2E). Finally, genes upregulated in Grh+ INPs were enriched for protein binding and system development (Fig1supp2D). Interestingly, we observed that the glial identity-promoting factor glial cell missing (gcm) and cell cycle inhibitor dacapo (dap) were upregulated in Ey^+^ INPs (Fig1G and supp1F). These observations support previous findings indicating that INPs begin producing glia cells instead of neurons during their later cell divisions, and that Ey is required for cell cycle exit (Baumgardt et al., 2014; Bayraktar & Doe, 2013; Ren, Awasaki, Wang, Huang, & Lee, 2018; Viktorin, Riebli, & Reichert, 2013). Unexpectedly, hamlet, a temporal switch gene required for the Grh-to-Ey transition (Eroglu et al., 2014), had high, but fluctuating expression levels during different temporal stages, suggesting that it might be required for correct progression of temporal patterning (Fig1G-I). This prompted us to focus on genes exhibiting similarly dynamic expression between INP subpopulations. Interestingly, odd-paired (opa), exhibited an expression pattern opposite of ham: high in D^+^ INPs, lower in Grh^+^ INPs and finally higher in Ey^+^ INPs (Fig1G-I). As Opa is a direct target of Osa (Eroglu et al., 2014) we investigated the potential role of Opa in regulating INP temporal patterning.

### Odd-paired (opa) is required for the progression of INP temporal patterning

Opa is a transcription factor containing five zinc finger domains and is essential for para-segmental subdivision of *Drosophila* embryos (Benedyk, Mullen, & DiNardo, 1994; Mizugishi, Aruga, Nakata, & Mikoshiba, 2001). During development, Opa ensures the timely activation of the transcription factors engrailed and wingless (Benedyk et al., 1994). To test if opa regulates INP temporal patterning, we depleted opa using RNAi expressed specifically in INPs with ermGal4. Opa knockdown slightly increased the total number of INPs (Dpn^+^ cells), but drastically increased the number of D^+^ INPs while decreasing the number of both Grh^+^ and Ey^+^ INPs (Fig2A-D). We confirmed this result by performing mosaic analysis with a repressible cell marker (MARCM) to create mosaic opa (-/-) mutant or control opa (+/+) GFP^+^ cell clones (T. Lee & Luo, 1999). Control clones were indistinguishable from WT, whereas opa mutant clones contained predominantly D^+^ INPs, at the expense of the other two temporal states (Fig2E-D). The RNAi and mosaic mutant analysis both indicate that loss of Opa causes a shift in INP temporal state identity such that the early generated D^+^ INPs are increased while the later generated Grh^+^ and Ey^+^ INPs are decreased.

**Figure 2.**
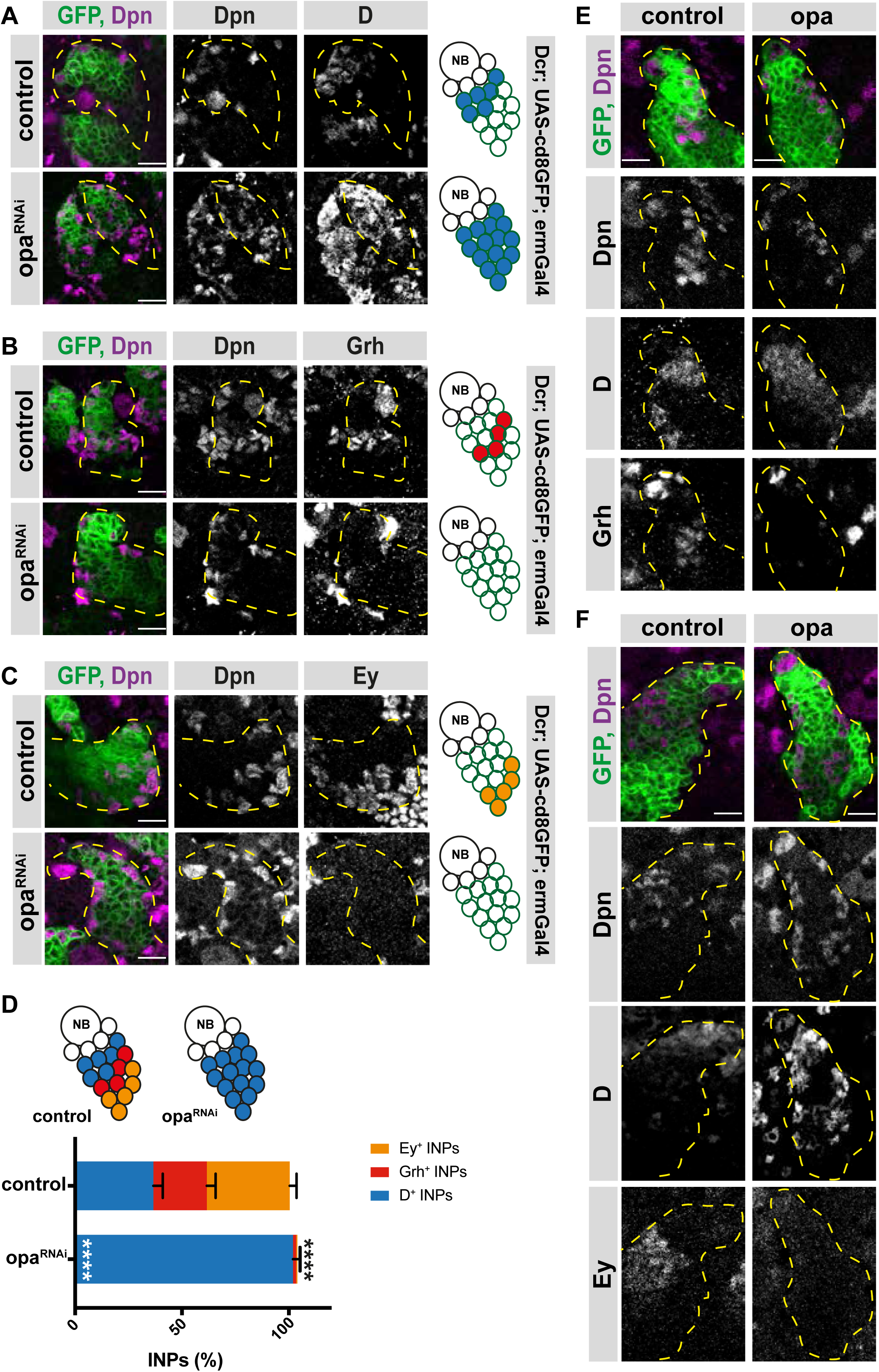
Opa is required for the progression of temporal patterning of INPs. (**A**) Close-up images of larval brains expressing RNAi against opa in INPs, stained for Dpn and D (induced with ermGal4, marked with membrane bound GFP). Lineages are outlined with yellow dashed line. (**B**) Close-up images of larval brains expressing RNAi against opa in INPs, stained for Dpn and Grh (induced with ermGal4, marked with membrane bound GFP). Lineages are outlined with yellow dashed line. (**C**) Close-up images of larval brains expressing RNAi against opa in INPs, stained for Dpn and Ey (induced with ermGal4, marked with membrane bound GFP). Lineages are outlined with yellow dashed line. (**D**) Quantification of INP numbers in different temporal stages identified by antibody staining of Dpn^+^, D^+^ cells, Dpn^+^, Grh^+^ cells, and Dpn^+^, Ey^+^ cells in control and opa knock-down brains, n=10, total INP numbers in control were normalized to 100%. Data represent mean ±SD, ***P<=0.001, Student’s t-test (D^+^ INPs control 12.44 ± 1.42 [n=10], opa RNAi 34.66 ± 1.02 [n=12], p<0.001; Grh^+^ INPs control 8.5 ± 1.32 [n=10], opa RNAi 0.5± 0.65 [n=12], p<0.001; Ey^+^ INPs control 13.2 ± 0.98 [n=10], opa RNAi 0.2 ± 0.4 [n=10], p<0.001). (**E**) Control and opa mutant MARCM clones marked by membrane-bound GFP, stained for Dpn, Grh and D after 120 hours of induction. Control clone has D^+^, Dpn^+^ INPs followed by Grh^+^ INPs while opa mutant clone has increased number of D^+^ INPs and decreased number of Grh^+^ INPs. (**F**) Control and opa mutant MARCM clones marked by membrane-bound GFP, stained for Dpn, D and Ey after 120 hours of induction. Opa mutant clone has higher number of D^+^ INPs and lower number of Ey^+^ INPs. Scale bar 10 μm in all images.

Finally, we tested if opa regulates processes upstream of temporal patterning during the stages of initial INP maturation with a type II specific driver line. When expressing opa RNAi specifically in type II NBs, we observed no effect on INP maturation, but immunofluorescent analysis of INPs for D, Grh and Ey expression showed the same phenotype as INPs depleted for opa (Fig2supp1A-C). Collectively, these data suggest that opa inhibits D expression. Furthermore, similar to hamlet, Opa appears to act as a temporal identity switch gene, controlling the transition from a D^+^ to a Grh^+^ state.

### Opa regulates the transition from early to late born neurons and is required for motor function

INP temporal patterning results in the production of different neuronal subtypes at distinct periods of neurogenesis. For instance, “young”, D^+^ INPs produce Brain-specific homeobox (Bsh^+^) neurons and “old”, Ey^+^ INPs produce Toy^+^ neurons (Bayraktar & Doe, 2013). Since the progression of INP temporal identity is disrupted in Opa-depleted INPs, we tested whether this disrupted identity affects the production of different types of neurons. Both INP-driven opa RNAi and opa-depleted MARCM clones displayed a significant increase in Brain-specific homeobox (Bsh)^+^ neurons, at the expense of Toy^+^ neurons (Fig3A-D). This result confirms that shifting the INP identity toward a D^+^ identity leads to a concomitant increase in the Bsh^+^ neurons produced by D^+^ INPs. Thus, altering the temporal identity progression of neural progenitors can alter the proportions of neuronal subtypes in the brain.

**Figure 3.**
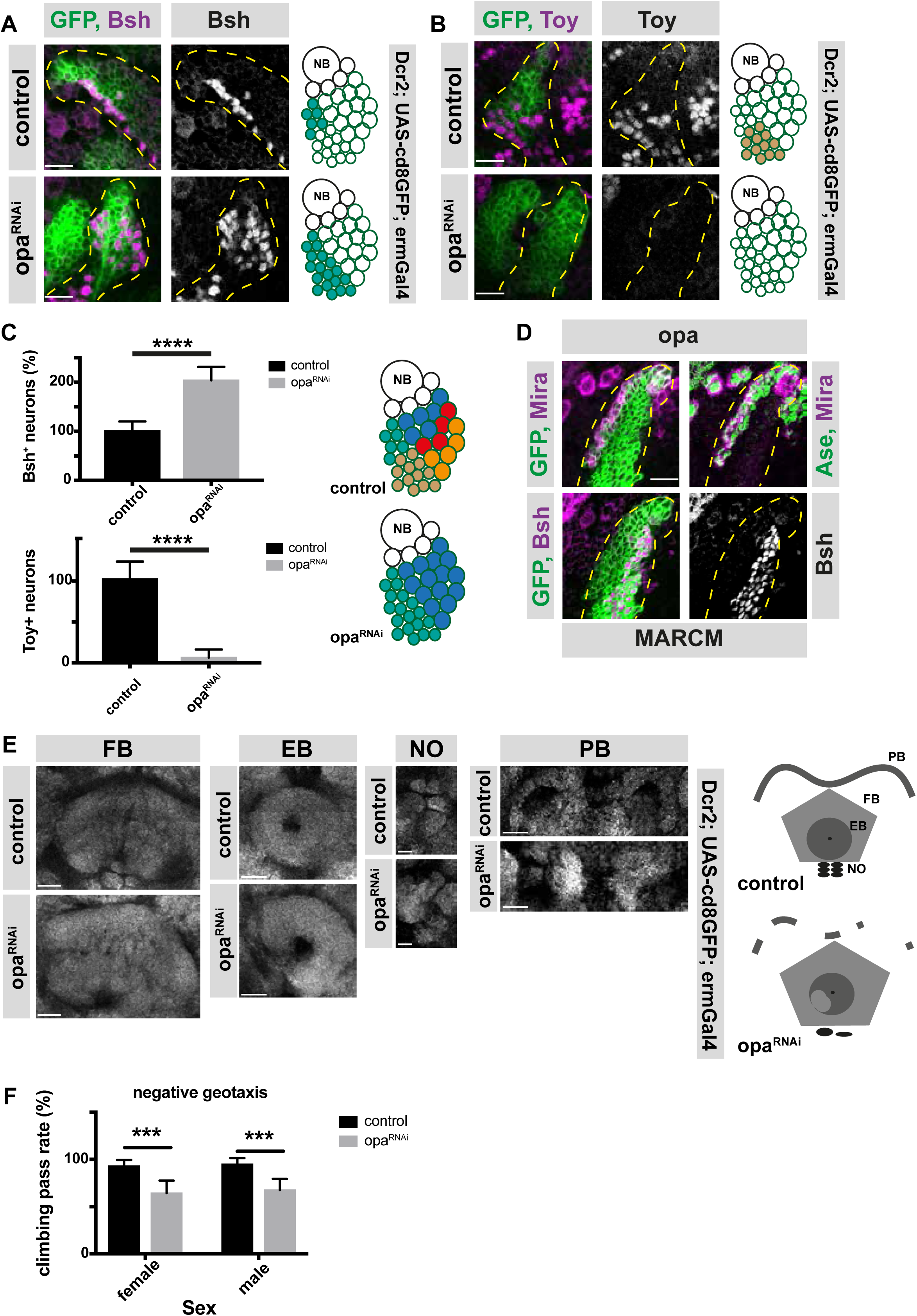
opa is an important factor for the generation of both early and late-born INP progeny and contributes to adult brain central complex. (**A-B**) Close-up images of larval brains expressing RNAi against opa in INPs, immunofluorescence for Bsh (**A**), and Toy (**B**) neuronal markers, scale bar 10 μm, lineages are outlined with yellow dashed line (induced with ermGal4, marked with membrane bound GFP). (**C**) Quantification of Bsh^+^ and Toy^+^ neurons in control and opa knock-down brains, n=11, total Bsh^+^ or Toy^+^ neuron numbers in control were normalized to 100%. Data represent mean ±SD, ***P<=0.001, Student’s t-test. (**D**) Opa mutant MARCM clone marked by membrane-bound GFP, stained with Mira, Ase, and Bsh antibodies after 120 hours of induction. The clone is marked with yellow dashed line. (**E**) Close-up images of adult central complex, composed of ellipsoid body (EB, scale bar 20 μm), fan-shaped body (FB, scale bar 20 μm), noduli (NO, scale bar 10 μm) and protocerebral bridge (PB, scale bar 20 μm) of control and opa knock-down brains, stained with Bruchpilot antibody (gray) (induced with ermGal4). (**F**) Negative geotaxis assay with control and opa RNAi expressing flies (induced with ermGal4, marked with membrane bound GFP). For each genotype n=10 replicates, each consisting of 10 adult female or male adults. Data are mean ± SD, ***P<0.001, Student’s t-test.

We next investigated whether altering the proportions of neuronal subtypes leads to a defect on brain morphology and function. The adult central complex (CCX) brain region relies on type II NB neurogenesis (Bayraktar, Boone, Drummond, & Doe, 2010; Izergina, Balmer, Bello, & Reichert, 2009). Opa-depletion in INPs caused major alterations in the gross morphology of the adult CCX. The fan-shaped body (FB) was enlarged, the noduli (NO) and ellipsoid body (EB) only partially formed, and the protocerebral bridge (PB) appeared fragmented (Fig3E). Since the CCX is required for adult motor functions (Callaerts et al., 2001; Young & Armstrong, 2010), we tested whether altered CCX morphology affected motor behavior. Compared to control flies, INP-driven *opa* RNAi caused impaired negative geotaxis performance (Fig3F). Thus, opa is a temporal switch gene required for neuronal subtype specification, which is required for the correct assembly and function of the adult central complex. Thus, the temporal identity specification of neural progenitors is crucial for proper neural cell complexity, and brain function.

### Dichaete and Opa are sequentially expressed in INPs

If opa is required for the D-to-grh transition, what is the molecular mechanism of this transitional regulation? To answer this question, we first confirmed that opa is indeed a target of Osa in type II NB lineages by analyzing opa protein expression within the NB lineage, and whether this expression is regulated by Osa. We generated healthy, homozygous, endogenously C-terminally tagged opa::V5 knock-in flies (Fig4supp1A). Through IF analysis of V5 tag expression, we observed that Opa is expressed throughout the type II lineage in INPs (marked with Dpn and Ase) and GMCs, but not in NBs (Dpn^+^) or immature INPs (Dpn^-^/Ase^-^ or Dpn^-^/Ase^+^ cells) (Fig4supp1A and B). The proper expression of opa depended on Osa, since Osa-knockdown in type II NBs resulted in a loss of Opa (Fig4supp1C and D). However, outside of the type II lineages, opa expression was readily detectable (Fig4supp1C), indicating that opa is directly activated by Osa in type II lineages specifically.

Since both D and opa are direct Osa targets, we next compared the expression pattern of D and opa (Fig4A). Without exception, D^+^/opa^-^ INPs appeared before D^+^/opa^+^ cell in the lineage (Fig4A). Since D expression precedes opa expression, it is possible that D activates opa. However, upon type II NB specific D knockdown, opa localization was unchanged (Fig4B). Interestingly, D knockdown alone also did not prevent later temporal stages, Grh and Ey, to appear (Bayraktar & Doe, 2013), suggesting that other factor(s) are required to maintain temporal identities in INPs. Since Osa-depleted type II NB lineages fail to initiate temporal patterning (Eroglu et al., 2014), we hypothesized that one of these unidentified factors could be a target of Osa that remains expressed in D-depleted INPs, such as opa. To test this hypothesis, we examined the epistatic genetic interactions between D and Opa. Double knock down of D and opa by type II NB-specific RNAi produced type II lineages containing fewer Dpn^+^/Ase^+^ INPs compared to controls (Fig4C-D). This result suggests that even though D and opa are Osa targets, two of them alone cannot fully account for Osa tumor suppressor role (Fig4D).

**Figure 4.**
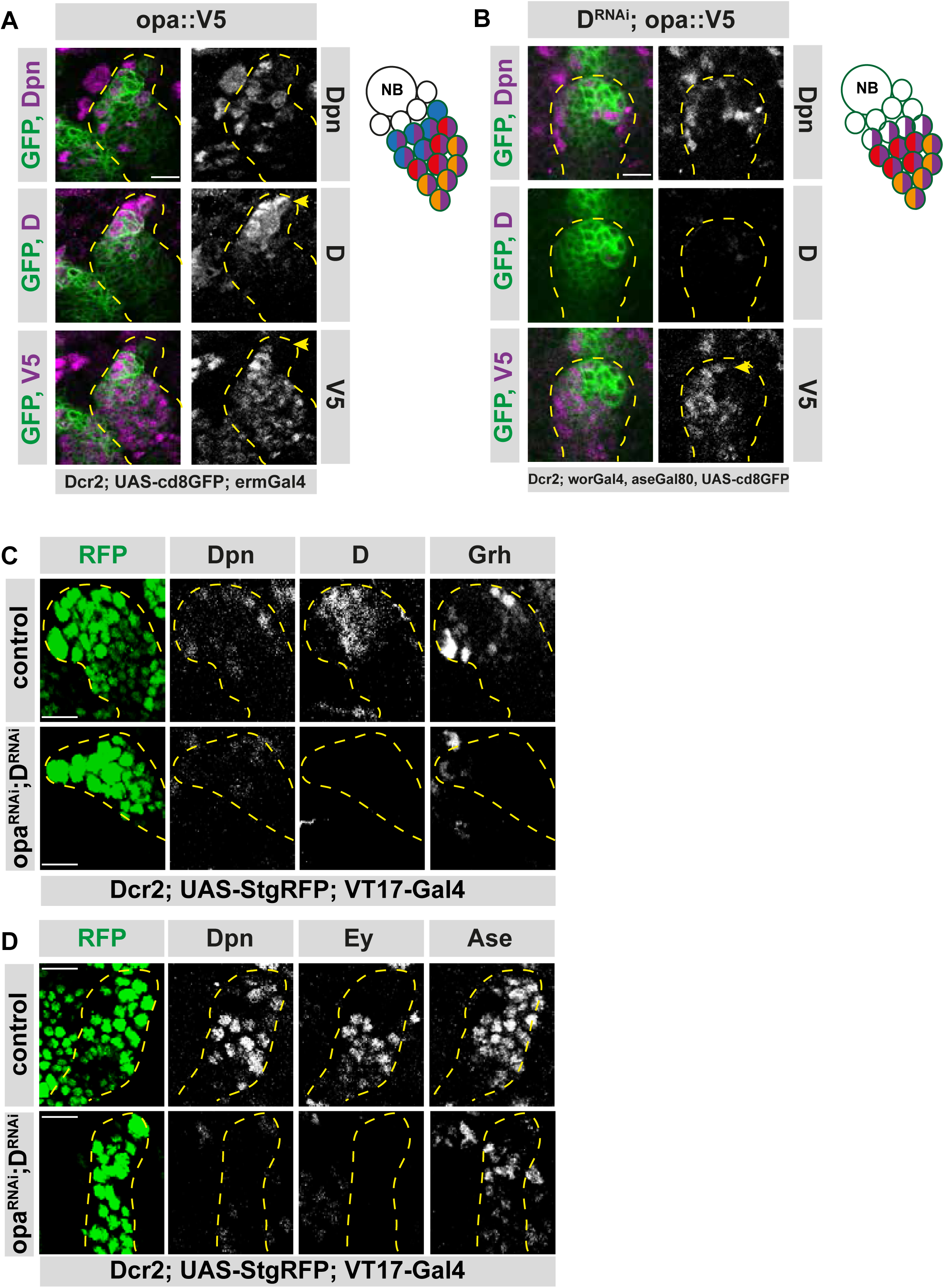
Osa initiates D expression before initiating Opa. (**A**) Close-up images of fly brains endogenously expressing V5-tagged opa in INPs, stained for V5, Dpn and D. D^+^, V5^-^ cell is marked with arrows, lineages are outlined with yellow dashed line, scale bar 10 μm, (induced with ermGal4, marked with membrane bound GFP). (**B**) Close-up images of fly brains endogenously expressing V5-tagged opa and RNAi for D in type II lineages, stained for V5, Dpn and D, lineages are outlined with yellow dashed line, scale bar 10 μm, (induced with worGal4, aseGal80, marked with membrane bound GFP). (**C-D**) Close up images of control versus opa and D double knock-down brains in type II lineages, stained with Dpn, D and Grh (**C**), or for Dpn, Ey and Ase (**C**) antibodies, lineages are outlined with yellow dashed lines, scale bar 10 μm, (induced with Dcr2; UAS-StgRFP; VT17-Gal4, marked with nuclear RFP).

Importantly, all known temporal identity markers on the remaining cells were absent, suggesting a complete loss of temporal identity in these INPs (Fig4C-D). However, since these cells also lost their INP identity due to lack of Dpn and Ase, they exhibit a different phenotype than Osa knockdown. Therefore, our data suggests that opa is required for the repression of D, the activation of Grh, and thus the initiation of temporal identity in INPs.

### Opa is an expression level-dependent repressor of D

If Opa suppresses D, one puzzling aspect of our data is the presence of double-positive D^+^opa^+^ INPs (Fig4A). To better understand this paradox, we overexpressed opa in type II NBs during a period before D is normally expressed. Overexpression of opa resulted in shorter lineages (Fig5supp1A-B), decreased total INP numbers (Fig5supp1A), and a loss of type II NBs (marked by Dpn or Mira) (Fig5supp1A-B). Co-expressing the apoptosis inhibitor p35 did not prevent NB loss or shortened lineages, suggesting that opa overexpression does not induce cell death, but causes premature differentiation instead (Fig5supp1C). Overexpressing opa in type II NB lineages caused complete loss of D^+^ INPs, but the few remaining INPs could still activate Grh and Ey (Fig5A-C), which is similar to D knockdown phenotype (Bayraktar & Doe, 2013).

To exclude that these could result from altered NB patterning, we next overexpressed opa in an INP-specific manner during a stage where D is normally expressed. Opa overexpression caused a decrease in D^+^ INPs (Fig5D-F), and a concomitant increase in both Grh^+^ and Ey^+^ INP populations (Fig5D-F). This result further indicates that Opa represses the early D^+^ temporal identity, but also activates later Grh^+^ temporal identity. Collectively, these results show that opa-mediated repression of D depends on Opa expression levels.

**Figure 5.**
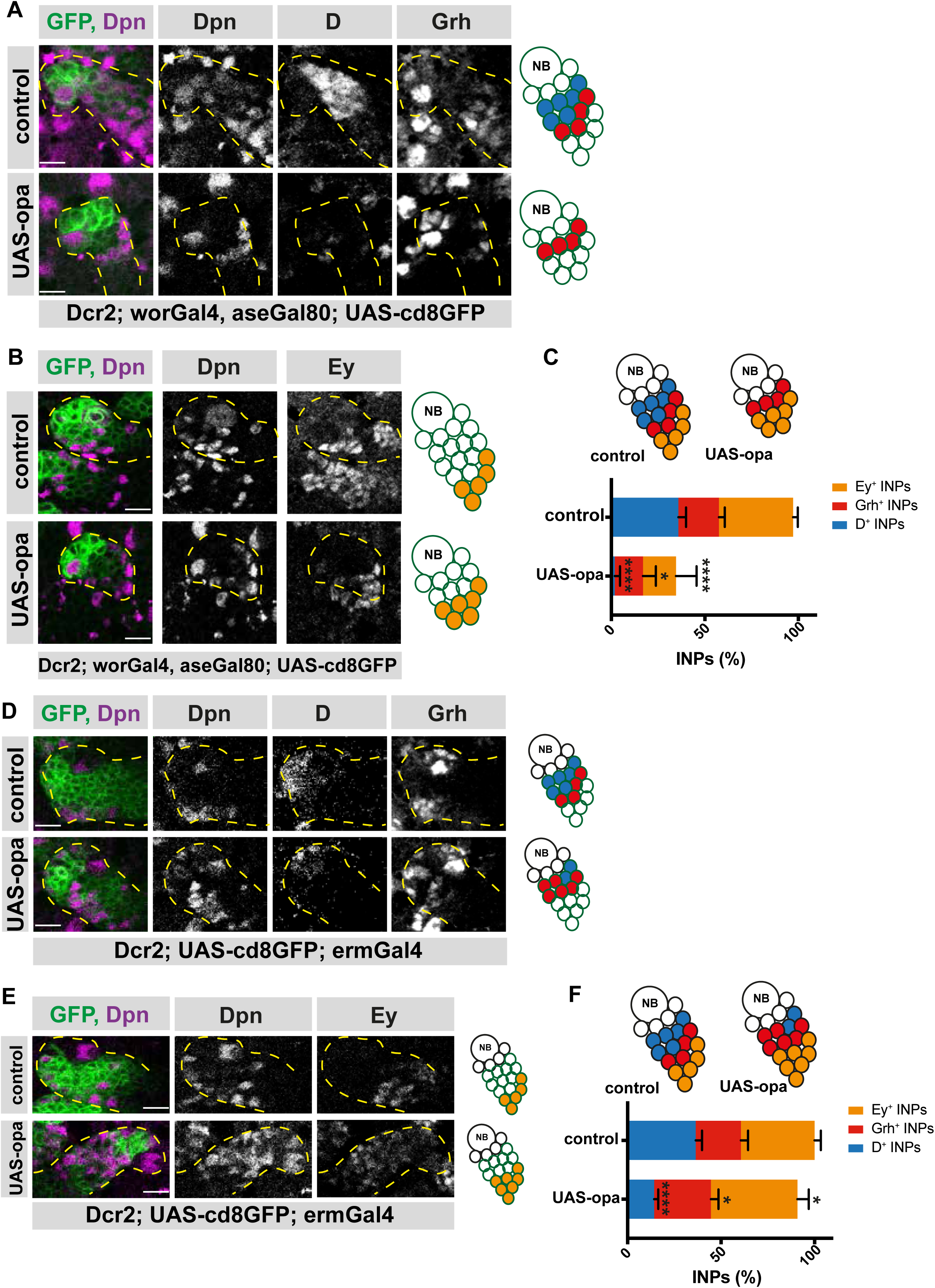
Opa overexpression results in the loss of D^+^ INPs. (**A**) Close-up images of control and opa overexpressing brains in type II lineages, stained for Dpn, D and Grh antibodies, lineages are outlined with yellow dashed lines, scale bar 10 μm, (induced with worGal4, aseGal80, marked with membrane bound GFP). Overexpression of opa in type II lineages causes the loss of D^+^ INPs. (**B**) Close-up images of control and opa overexpressing brains in type II lineages, stained for Dpn, and Ey antibodies, lineages are outlined with yellow dashed lines, scale bar 10 μm, (induced with worGal4, aseGal80, marked with membrane bound GFP). (**C**) Quantification of D^+^, Grh^+^ and Ey^+^ INPs in control and opa overexpressing brains, n=10, total INP numbers in control were normalized to 100%. Data represent mean ±SD, P<=0.05, ***P<=0.001, Student’s t-test (D^+^ INPs control 12.18 ± 1.33 [n=10], opa GOF 0.4 ± 0.6 [n=10], p<0.001; Grh^+^ INPs control 7.38 ± 1 [n=10], opa GOF 5.12 ± 2.20 [n=10], p<0.05; Ey^+^ INPs control 13.5 ± 0.76 [n=10], opa GOF 6 ± 3.5 [n=10], p<0.001). (**D**) Close-up images of control and opa overexpressing brains in INPs, stained for Dpn, and Ey, lineages are outlined with yellow dashed lines, scale bar 10 μm, (induced with ermGal4, marked with membrane bound GFP). (**E**) Close-up images of control and opa overexpressing brains in INPs, stained for Dpn, D and Grh, lineages are outlined with yellow dashed lines, scale bar 10 μm, (induced with ermGal4, marked with membrane bound GFP). (**F**) Quantification of D^+^, Grh^+^ and Ey^+^ INPs in control and opa overexpressing brains, n=5, total INP numbers in control were normalized to 100%. Data represent mean ±SD, *P<=0.05, ***P<0.001, Student’s t-test (D^+^ INPs control 12.4 ± 1.01 [n=5], opa GOF 4.83 ± 0.68 [n=5], p<0.001; Grh^+^ INPs control 8.2 ± 1.16 [n=5], opa GOF 10.33 ± 1.24 [n=5], p<0.05; Ey^+^ INPs control 13.4 ± 1.01 [n=5], opa GOF 15.71 ± 1.9 [n=5], p<0.05).

### Opa and ham together control the correct representation of each temporal identity

Having established an interaction between opa and D, we next wondered if opa and ham, two temporal switch genes, can recapitulate the Osa loss-of-function phenotype, a more upstream regulator of lineage progression in type II NBs. Osa knock-down causes INPs to revert back to the NB-state due to a failure to initiate temporal patterning, while single depletion of opa or ham leads to either an increase in D^+^ or Grh^+^ cells, respectively (Fig2, (Eroglu et al., 2014)). Co-expressing opa RNAi with ham shmiR in an INP-specific manner caused supernumerary Dpn^+^, Ase^+^ INPs (Fig6supp1A). In addition, the number of D^+^/Dpn^+^ and Grh^+^/Dpn^+^ INPs were also increased, which is in contrast to single depletion of opa or ham (Fig6A-B, Fig2, (Eroglu et al., 2014)). Thus, opa and ham loss-of-function phenotypes are additive. Importantly, despite inducing over-proliferation of mature INPs (Ase^+^/Dpn^+^), depleting both opa and ham in type II NBs could not recapitulate the Osa loss-of-function phenotype because imINPs could mature and express Ase, and therefore did not revert into ectopic NBs (Fig6supp1B). This suggests that Osa regulates temporal patterning in two levels: initiation by D activation, and progression by opa and ham.

## Discussion

Temporal patterning is a phenomenon where NSCs alter the fate of their progeny chronologically. Understanding how temporal patterning is regulated is crucial to understanding how the cellular complexity of the brain develops. Here, we present a novel, FACS-based approach that enabled us to isolate distinct temporal states of neural progenitors with very high purity from Drosophila larvae. This allowed us to study the transitions between different temporal identity states. We identified odd-paired (opa), a transcription factor that is required for INP temporal patterning. By studying the role of this factor in temporal patterning, we propose a novel model for the regulation of temporal patterning in *Drosophila* neural stem cells.

We establish two different roles of the SWI/SNF complex subunit, Osa, in regulating INP temporal patterning. Initially, osa initiates temporal patterning by activating the transcription factor D. Subsequently, osa regulates the progression of temporal patterning by activating opa and ham, which in turn downregulate D and Grh, respectively (Fig6C). The concerted, but complementary action of opa and ham ensures temporal identity progression by promoting the transition between temporal stages. For instance, opa regulates the transition from D to Grh, while ham regulates the transition from Grh to Ey. We propose that opa achieves this by repressing D and activating grh, as indicated by the lack of temporal patterning in D and opa-depleted INPs (Fig4C-D, Fig6C). Loss of opa or ham causes INPs to lose their temporal identity and overproliferate. Moreover, we propose that D and opa activate Grh expression against the presence of ham, which represses Grh expression. As D and opa levels decrease as INPs age and become Grh positive, ham is capable of repressing Grh later on in temporal patterning (Fig6C). This explains how opa and ham act only during specific stages even though they are expressed throughout the entire lineage

**Figure 6.**
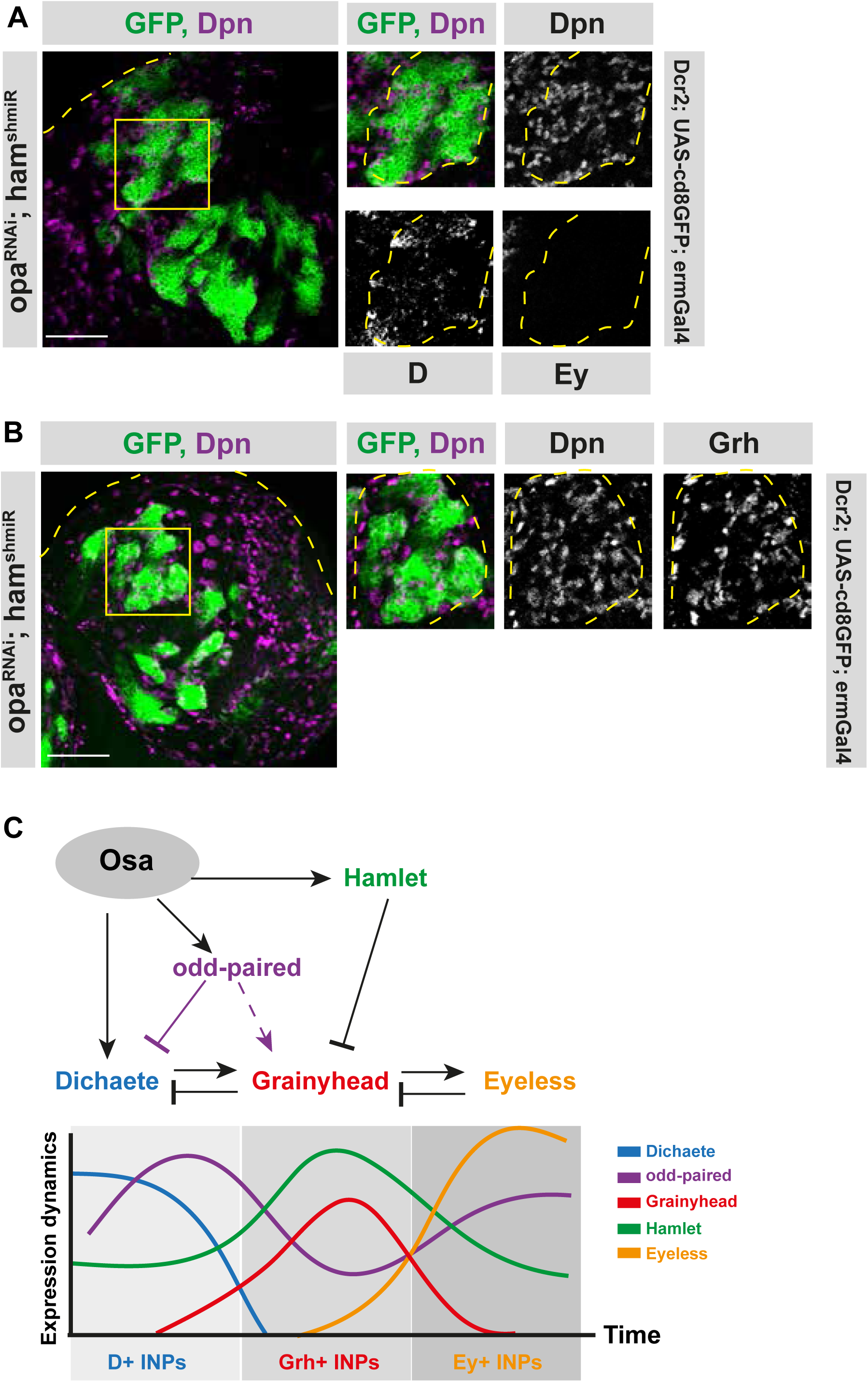
Opa and hamlet are required for INP temporal patterning and correct lineage progression. (**A-B**) Overview images of brain lobes expressing RNAi against opa and ham in INPs and their close-up images (marked with yellow squares), stained for Dpn, D and Ey (**A**), or Dpn and Grh (**B**) antibodies, lineages and lobes are outlined with yellow dashed lines, scale bar 50 μm for brain lobes, 10μm for zoomed images, (induced with ermGal4, marked with membrane bound GFP). (**C**) Model depicting the genetic interactions between temporal switch genes (opa and hamlet), and temporal identity genes (D, Grh, and Ey).

An open question pertains to the fact that the double knock-down of opa and ham, as well as that of D and opa, failed to recapitulate the Osa phenotype. Even though opa and ham RNAi caused massive overproliferation in type II lineages, we could not detect any Dpn^+^ Ase^-^ ectopic NB-like cells (as occurs in Osa mutant clones, (Eroglu et al., 2014)). We propose that this is caused by D expression which is still induced even upon opa/ham double knockdown, but not upon Osa knock-down where D expression fails to be initiated. Thus, the initiation of the first temporal identity state may block the reversion of INPs to a NB-state. In the future, it will be important to understand the exact mechanisms of how opa regulates temporal patterning.

We further demonstrate that Osa initiates D expression earlier than opa expression. Osa is a subunit of SWI/SNF chromatin remodeling complex, and it guides the complex to specific loci throughout the genome, such as the TSS of both D and opa. The differences in timing of D and opa expression may be explained by separate factors involved in their activation. Previous work suggests that the transcription factor earmuff may activate (Janssens et al., 2014; 2017). However, it remains unknown which factor activates opa expression. One possibility is that the cell cycle activates opa (Isshiki et al., 2001), since its expression begins in mINPs, a dividing cell unlike imINPs, which are in cell cycle arrest.

We propose that balanced expression levels of D and opa regulates the timing of transitions between temporal identity states. Indeed, Osa initiates D and opa, the repressor of D, at slightly different times, which could allow a time window for D to be expressed, perform its function, then become repressed again by opa. Deregulating this pattern, for example by overexpressing opa in the earliest INP stage, results in a false start of temporal patterning and premature differentiation. This elegant set of genetic interactions resembles that of an incoherent feedforward loop (FFL) (Dongsan Kim, Kwon, & Cho, 2008; Mangan & Alon, 2003). In such a network, pathways have opposing roles. For instance, Osa promotes both the expression and repression of D. Similar examples can be observed in other organisms, such as in the galactose network of E.coli, where the transcriptional activator CRP activates galS and galE, while galS also represses galE (Shen-Orr, Milo, Mangan, & Alon, 2002). In Drosophila SOP determination, miR-7, together with Atonal also forms an incoherent FFL (X. Li, Cassidy, Reinke, Fischboeck, & Carthew, 2009). Furthermore, mammals apply a similar mechanism in the c-Myc/E2F1 regulatory system (O’Donnell, Wentzel, Zeller, Dang, & Mendell, 2005).

The vertebrate homologues of opa consist of the Zinc-finger protein of the cerebellum (ZIC) family, which are suggested to regulate the transcriptional activity of target genes, and to have a role in CNS development (Elms et al., 2004; Elms, Siggers, Napper, Greenfield, & Arkell, 2003; Gaston-Massuet, Henderson, Greene, & Copp, 2005; Inoue et al., 2004; Inoue, Ota, Ogawa, Mikoshiba, & Aruga, 2007). In mice, during embryonic cortical development, ZIC family proteins regulate the proliferation of meningeal cells, which are required for normal cortical development (Inoue, Ogawa, Mikoshiba, & Aruga, 2008). In addition, another member of the ZIC family, Zic1, is a Brn2 target, which itself controls the transition from early-to-mid neurogenesis in the mouse cortex (Urban et al., 2015). Along with these lines, it has been shown that ZIC family is important in brain development in zebrafish (Maurus & Harris, 2009; Sanek & Grinblat, 2008). Furthermore, the role of ZIC has been implicated in variety of brain malformations and/or diseases (Aruga, Nozaki, Hatayama, Odaka, & Yokota, 2010; Blank et al., 2011; Hatayama et al., 2011). These data provide mere glimpses into the roles of ZIC family proteins in neuronal fate decisions in mammals, and our study offers an important entry point to start understanding these remarkable proteins.

Our findings provide a novel regulatory network model controlling temporal patterning, which may occur in all metazoans, including humans. In contrast to existing cascade models, we instead show that temporal patterning is a highly coordinated ensemble that allows regulation on additional levels than was previously appreciated to ensure a perfectly balanced generation of different neuron/glial cell types. Together, our results demonstrate that *Drosophila* is a powerful system to dissect the genetic mechanisms underlying the temporal patterning of neural stem cells and how the disruption of such mechanisms impacts brain development and behavior.

## Materials and Methods

### Fly strains, RNAi, and clonal analysis

The following *Drosophila* stocks were used: UAS-*opa*^RNAi^ (VDRC, TID: 101531), UAS-mcherry^shmiR^ (BL35785), UAS-D^RNAi^ (VDRC, TID: 49549, 107194), UAS-osa^RNAi^ (VDRC, TID: 7810), UAS-ham^shmiR^ (BL32470), UAS-osa^shmiR^ (Eroglu et al., 2014), UAS-p35, UAS-opa (H. Lee, Stultz, & Hursh, 2007), PBac{grh-GFP.FPTB}VK00033 (BL42272), PBac{EyGFP.FPTB}VK00033 (BL42271), D::GFP (generated in this study), opa::V5 (generated in this study). GAL4 driver lines used: UAS-cd8::tdTomato; *erm*Gal4, UAS-cd8::GFP; *erm*Gal4 (Pfeiffer et al., 2008; Weng, Golden, & Lee, 2010), UAS-*dcr2*; *wor*Gal4, *ase*Gal80; UAS-cd8::GFP (Neumüller et al., 2011), UAS-*dcr2*; UAS-cd8::GFP; VT17-Gal4 (VDRC, TID: 212057, discarded). Mutant fly strains used for clonal analysis were FRT82B, *opa*^7^ (H. Lee et al., 2007). Clones were generated by Flippase (FLP)/FLP recombination target (FRT)-mediated mitotic recombination, using the *elav*Gal4 (C155) (T. Lee & Luo, 1999). Larvae were heat shocked for 90 min at 37°C and dissected as third-instar wandering larvae (120 hours). RNAi crosses were set up and reared at 29°C, and five days later, third-instar wandering larvae were dissected. *w*^*118*^ was used as control for comparison with RNAi lines, whereas UAS-mcherry^shmiR^ was used as control for comparison with shmiR lines, and experiments involving UAS-transgenes.

### Generation of opa::V5 and D::GFP flies

For both genes, the guides were cloned as overlapping oligos into linearized pU6-BbsI-chiRNA (Addgene 45946, (Gratz et al., 2013)) and injected at 100 ng/μl into actCas9 flies (BL 54590, (Port, Chen, Lee, & Bullock, 2014)). Donors (either oligos or plasmid) were co-injected at 250 ng/μl. For opa, donors were Ultramer Oligos from IDT with around 60nt homology arms on either side. For D, homology arms were 800bp and 900bp long. Donor plasmid contained GFP, V5, 3xFlag, and dsRed. They were screened for dsRed eyes and then, the selection cassette was removed with hsCre (BL 851).

*opa* gRNA GATGCATCCCGGCGCAGCGA

*opa* donor

GAACCCGCTGAACCATTTCGGACACCATCACCACCACCACCACCTGATGCATCCCGGCGCgGCaACcGCGTATggtaagcctatacctaaccctcttcttggTCTAGAtagcacgTGAGAGTGGGAGAACTGGTGGCCCGAGGAGGCGCCACCGCCGGCCGCCCAACCGA

*D* gRNA GTGCTCTATTAGAGTGGAGT

### Negative geotaxis assay

Negative geotaxis assay was used as described before (Ali, Escala, Ruan, & Zhai, 2011), where the percentage of flies passing the 8.5 cm mark in 10 seconds was assessed. For each genotype and gender, 10 two-day old adult flies in 10 biological replicates were measured and for each replicate, 10 measurements were performed with 1 min rest period in between.

### Immunohistochemistry and Antibodies

Larval or adult brains were dissected in 1X PBS, and then fixed for 20 min at room temperature (RT) in 5% paraformaldehyde in PBS and washed once with 0.1% TritonX in PBS (PBST). The brains were incubated for 1 hour at RT with blocking solution (5% normal goat serum or 1% BSA in PBST). Blocking was followed by overnight incubation at 4°C with primary antibodies in blocking solution. Then, the brains were washed three times with PBST, and incubated for 1 hour at RT with secondary antibodies (1:500, goat Alexa Fluor®, Invitrogen) in blocking solution. After secondary antibody, brains were washed three times with PBST, and mounted in Vectashield Antifade Mounting Medium (Vector Labs).

Antibodies used in this study were: guinea pig anti-Deadpan (1:1000, (Eroglu et al., 2014)), rat anti-Asense (1:500, (Eroglu et al., 2014)), guinea pig anti-Miranda (1:500, (Eroglu et al., 2014)), rat anti-Grh (1:1,000; (Baumgardt, Karlsson, Terriente, Díaz-Benjumea, & Thor, 2009)); rabbit anti-D (1:1,000; gift from Steve Russell); mouse anti-Ey (1:10; DSHB); guinea pig anti-Toy (gift from Uwe Walldorf), guinea pig anti-Bsh (gift from Makoto Sato), mouse anti-Bruchpilot nc82 (1:10, DSHB). Throughout the paper, for every quantification, dorsomedial 2 and 3 type II NB lineages (DM2 and 3) were considered.

### In vitro Immunofluorescence

FACS-sorted cells from ∼300 larval brains (UAS-cd8::tdTomato, *erm*Gal4) or their unsorted control matches were plated on cover glass (Labtek II Chambered Coverglass, 8-well, 155409, Thermo Fisher Scientific) into Schneider’s medium (Homem, Reichardt, Berger, Lendl, & Knoblich, 2013). The dishes were placed onto ice and cells were incubated for 1 hour to settle down. Cells were then fixed with 5% PFA in PBS at RT and washed three times with 0.1% PBST. After washes, cells were incubated for 1 hour at RT with blocking solution (5% normal goat serum in 0.1% PBST). The cells were then incubated overnight at 4°C with primary antibodies in blocking solution, which was followed by three washes with 0.1% PBST, and secondary antibody (1:500, goat Alexa Fluor®, Invitrogen) incubation for 1 hour at RT. Cells were again washed three times with 0.1% PBST, and then mounted in in Vectashield Antifade Mounting Medium with Dapi (Vector Labs).

### Microscopy

Confocal images were acquired with Zeiss LSM 780 confocal microscopes.

### Statistics

Statistical analyses were performed with GraphPad Prism 7. Unpaired two-tailed Student’s *t*-test was used to assess statistical significance between two genotypes. Experiments were not randomized, and investigator was not blinded. Sample sizes for experiments were estimated on previous experience with similar setup which showed significance, thus, no statistical method was used to determine sample size.

### Cell dissociation and FACS

Cell dissociation and FACS were performed as previously described with minor changes (Berger et al., 2012; Harzer, Berger, Conder, Schmauss, & Knoblich, 2013). UAS-cd8::tdTomato; *erm*Gal4 driver line was used to induce expression of membrane bound tdTomato in INPs. In addition to the driver lines, temporal identity factors were tagged with GFP. Flies expressing both fluorophores were dissected at L3 stage, and then dissociated into single cell suspension. Decreasing levels of tdTomato were observed in differentiated cells due to lack of driver line expression. Thus, biggest cells with highest tdTomato expression and highest GFP expression were sorted. For RNA isolation, cells were sorted directly in TRIzol LS (10296010, Invitrogen), while for cell staining, they were sorted on coated glass-bottomed dishes and stained as previously described (Berger et al., 2012).

### RNA isolation, cDNA synthesis and qPCR

RNA was isolated using TRIzol LS reagent (10296010, Invitrogen) from FACS sorted cells. Then RNA samples were used as template for first-strand cDNA synthesis with random hexamer primers (SuperScript®III, Invitrogen). qPCR was done using Bio-Rad IQ SYBR Greeen Supermix on a Bio-Rad CFX96 cycler. Expression of each gene was normalized to Act5c, and relative levels were calculated using the 2^-ΔΔCT^ method (Livak & Schmittgen, 2001). Primer used were:

*act5c* AGTGGTGGAAGTTTGGAGTG, GATAATGATGATGGTGTGCAGG

*D* ATGGGTCAACAGAAGTTGGGAG, GTATGGCGGTAGTTGATGGAATG

*grh* TCCCCTGCTTATGCTATGACCT, TACGGCTAGAGTTCGTGCAGA

*ey* TCGTCCGCTAACACCATGA, TGCTCAAATCGCCAGTCTGT

*ham* ATAGATCCTTTGGCCAGCAGAC, AGTACTCCTCCCTTTCGGCAAT

*opa* CTGAACCATTTCGGACACCATC, CCAGTTCTCCCACTCTCAATAC

### RNA sequencing – DigiTAG

For each experiment 6000-7000 FACS-sorted D^+^, Grh^+^ or Ey^+^ INPs were isolated by TRIzol purification. Three replicates from each temporal state were analyzed. RNA samples were reverse transcribe into first-strand cDNA using SuperScript®III Reverse Transcriptase (Invitrogen) with oligo-(dT)2-primers. Then the second-strand cDNA were generated. It was followed by library preparation with Nextera DNA Library Preparation Kit (Illumina) as previously described (Landskron et al., 2018; Wissel et al., 2018). Libraries were purified with Agencourt AMPure XP beads. Purified libraries were then subjected to 50 base pair Illumina single-end sequencing on a Hiseq2000 platform.

### Transcriptome data analysis

#### Alignment

Unstranded reads were screened for ribosomal RNA by aligning with BWA (v0.7.12; (H. Li & Durbin, 2009)) against known rRNA sequences (RefSeq). The rRNA subtracted reads were aligned with TopHat (v2.1.1; (Daehwan Kim et al., 2013)) against the Drosophila genome (FlyBase r6.12). Introns between 20 and 150,000 bp are allowed, which is based on FlyBase statistics. Microexon-search was enabled. Additionally, a gene model was provided as GTF (FlyBase r6.12).

#### Deduplication

Reads arising from duplication events are marked as such in the alignment (SAM/BAM files) as follows. The different tags are counted at each genomic position. Thereafter, the diversity of tags at each position is examined. First, tags are sorted descending by their count. If several tags have the same occurrence, they are further sorted alphanumerically. Reads sharing the same tag are sorted by the mean PHRED quality. Again, if several reads have the same quality, they are further sorted alphanumerically. Now the tags are cycled through by their counts. Within one tag, the read with the highest mean PHRED quality is the unique correct read and all subsequent reads with the same tag are marked as duplicates. Furthermore, all reads that have tags with one mismatch difference compared the pool of valid read tags are also marked as duplicates.

#### Summarization

Small nuclear RNA, rRNA, tRNA, small nucleolar RNA, and pseudogenes are masked from the GTF (FlyBase r6.12) with subtractBed from bedtools (v2.26.0; (Quinlan & Hall, 2010)). The aligned reads were counted with HTSeq (v0.6.1; intersection-nonempty), and genes were subjected to differential expression analysis with DESeq2 (v1.12.4; (Love, Huber, & Anders, 2014)).

### Hierarchical clustering analysis

Genes are filtered by the indicated log2fc and an adjusted P value < 0.05 in at least one pairwise comparison. In addition, a minimal expression of 10 RPM in at least one condition was required. The tree cut into four clusters (different cluster numbers were tested; (Kolde, Package, 2015, 202AD)). GO analysis was performed with FlyMine (Lyne et al., 2007), Holm-Bonferroni correction with max p-value 0.05 was used. Biological process and molecular function were the ontologies.

### Accession numbers

The Gene Expression Omnibus accession number for the RNA-sequencing data reported in this paper is GSE127516.

### GO-term analysis

Gene Ontology (GO) enrichment analysis were performed on www.flymine.org/ with Holm-Bonferroni correction with max p-value 0.05. Biological process and molecular function were the ontologies.

## Acknowledgement

We thank all Knoblich lab members for support and discussions, Francois Bonnay, Tom Kruitwagen and Joshua A. Bagley for comments on the manuscript, Peter Duchek, Joseph Gokcezade, Elke Kleiner, the IMP/IMBA Biooptics Facility and the Next Generation Sequencing Unit of the Vienna Biocenter Core Facilities (VBCF) for assistance and Makoto Sato, Uwe Walldorf, Stefan Thor and Steve Russel for sharing reagents, and the Harvard TRiP collection, the Bloomington Drosophila stock center and the Vienna Drosophila Resource Center (VDRC) for reagents.

## Author contributions

M.D.A. conceptualized, designed, performed, interpreted experiments and wrote the manuscript. E.E. conceptualized and performed experiments, T. R. B. conducted all bioinformatic analyses. J.A.K. supervised the project, planned and interpreted experiments, and wrote the manuscript.

---

**Figure1 supplemet1.**
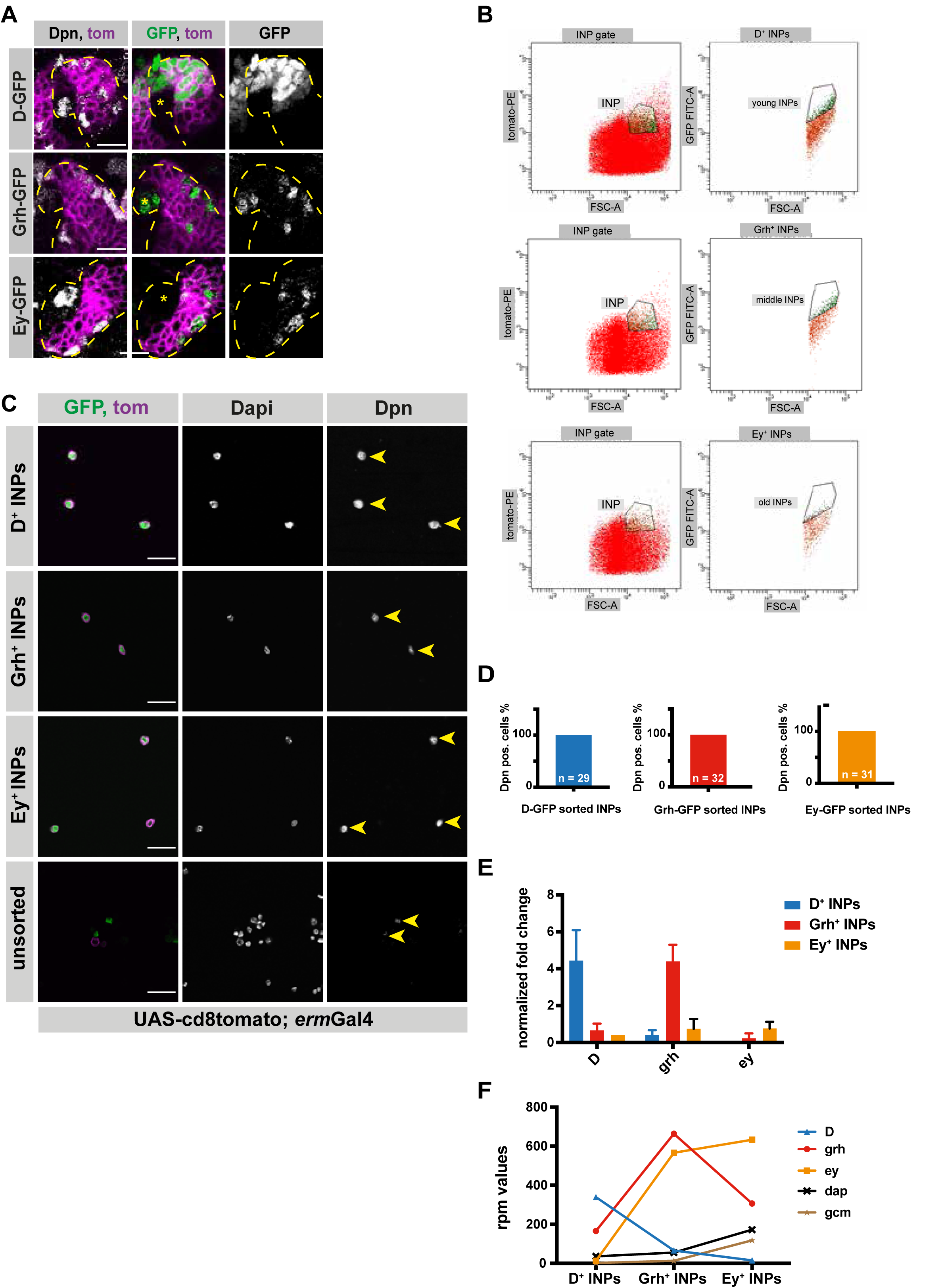
INPs can be FACS-sorted depending on their temporal identity. (**A**) Close-up images of larval brains expressing tdtomato in INP-specific manner (UAS-cd8tomato; ermGal4, shown in magenta) and temporal identity markers, Dichaete, Grainyhead and Eyeless tagged with GFP, stained for Dpn (gray). Type II lineages are outlined with yellow dashed lines, scale bar 10 μm, (induced with ermGal4, marked with membrane bound tdTomato). (**B**) Representative FACS plots of sorted INP temporal stages. The population with highest tomato and GFP expression (vertical axis) versus biggest cell size (horizontal axis) was sorted to obtain D^+^, Grh^+^, and Ey^+^ INPs. (**C**) D-, Grh-and Ey-GFP FACS-sorted cells and their unsorted control are stained for Dpn (gray), GFP-tagging temporal identity factors (in green, D or, Grh or Ey), tdTomato tagging the membrane of INPs (magenta), Dapi (gray), scale bar 20 μm, yellow arrowheads mark Dpn positive cells, (induced with ermGal4, marked with membrane bound tdTomato). (**D**) Graph showing the percentages of Dpn positive cells, n is the number of the cells analyzed. (**E**) qPCR analysis of temporal identity expression levels in FACS-sorted D^+^, Grh^+^ and Ey^+^ INPs. Data are mean ± SD, n=3, genes were normalized to Act5c, and then the average expression levels, Delta-Delta Ct method is used. (**F**) Graph showing the rpm levels of marker genes between different INP temporal stages.

**Figure1 supplement2.**
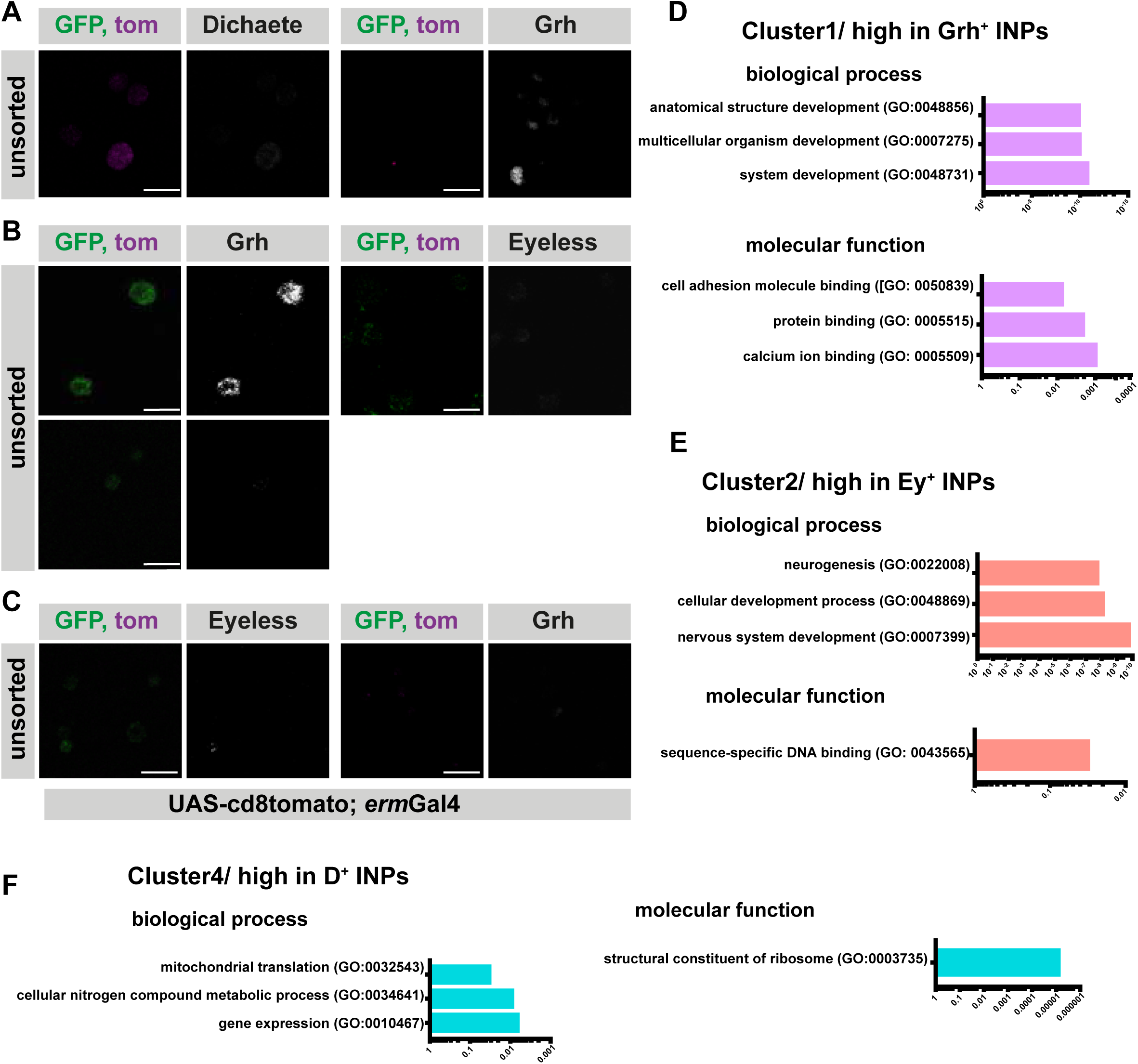
Temporally sorted INPs are pure populations. (**A-C**) Immunofluorescence of unsorted controls for Fig1C-E, scale bar 10 μm, (induced with ermGal4, marked with membrane bound tdTomato). (**D-F**) GO-term analysis of each cluster found in hierarchical clustering analysis for biological process, and molecular function. The graphs are color-coded with their respective clusters, the top 3 hits were shown if applicable. Cluster 2 doesn’t have any GO-term enrichment.

**Figure2 supplement1.**
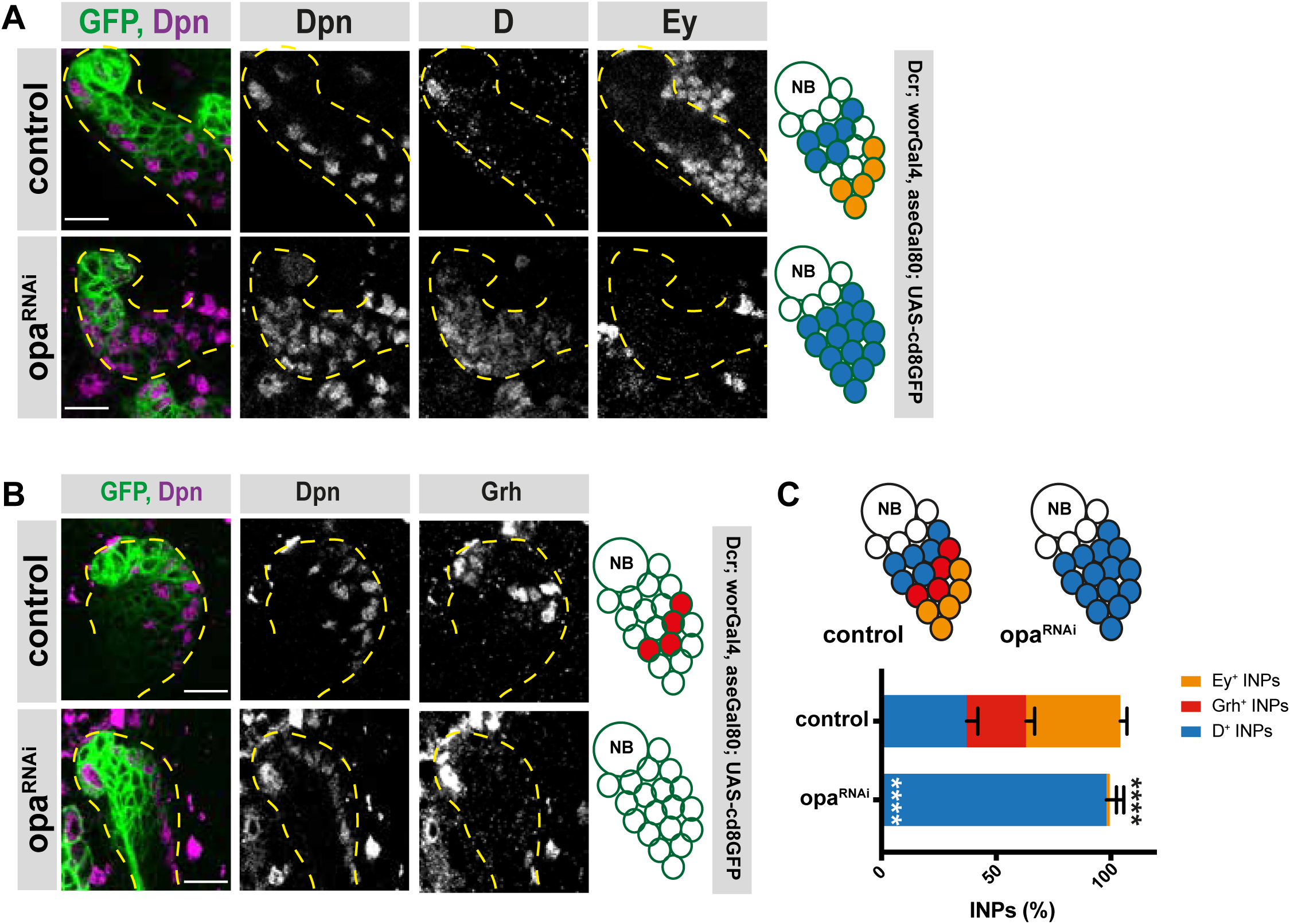
Opa regulates the transition from D-to-grh. (**A-B**) Close-up images of larval brains expressing RNAi against opa in type II lineages, stained for Dpn, D and Ey (**A**), and Grh (**B**), lineages are outlined with yellow dashed line, scale bar 10 μm (induced with worGal4, aseGal80, marked with membrane bound GFP). (**C**) Quantification of Dpn^+^, D^+^, and Dpn^+^, Grh^+^, and Dpn^+^, Ey^+^ INPs in control and opa knock-down brains, n=5, total INP numbers in control were normalized to 100%. Data represent mean ±SD, ***P<=0.001, Student’s t-test. (D^+^ INPs control 12.6 ± 1.5 [n=5], opa RNAi 33.3 ± 2.35 [n=6], p<0.001; Grh^+^ INPs control 8.8 ± 1.6 [n=5], opa RNAi 0 [n=6], p<0.001; Ey^+^ INPs control 14 ± 0.89 [n=5], opa RNAi 0.5 ± 0.86 [n=5], p<0.001).

**Figure4 supplement1.**
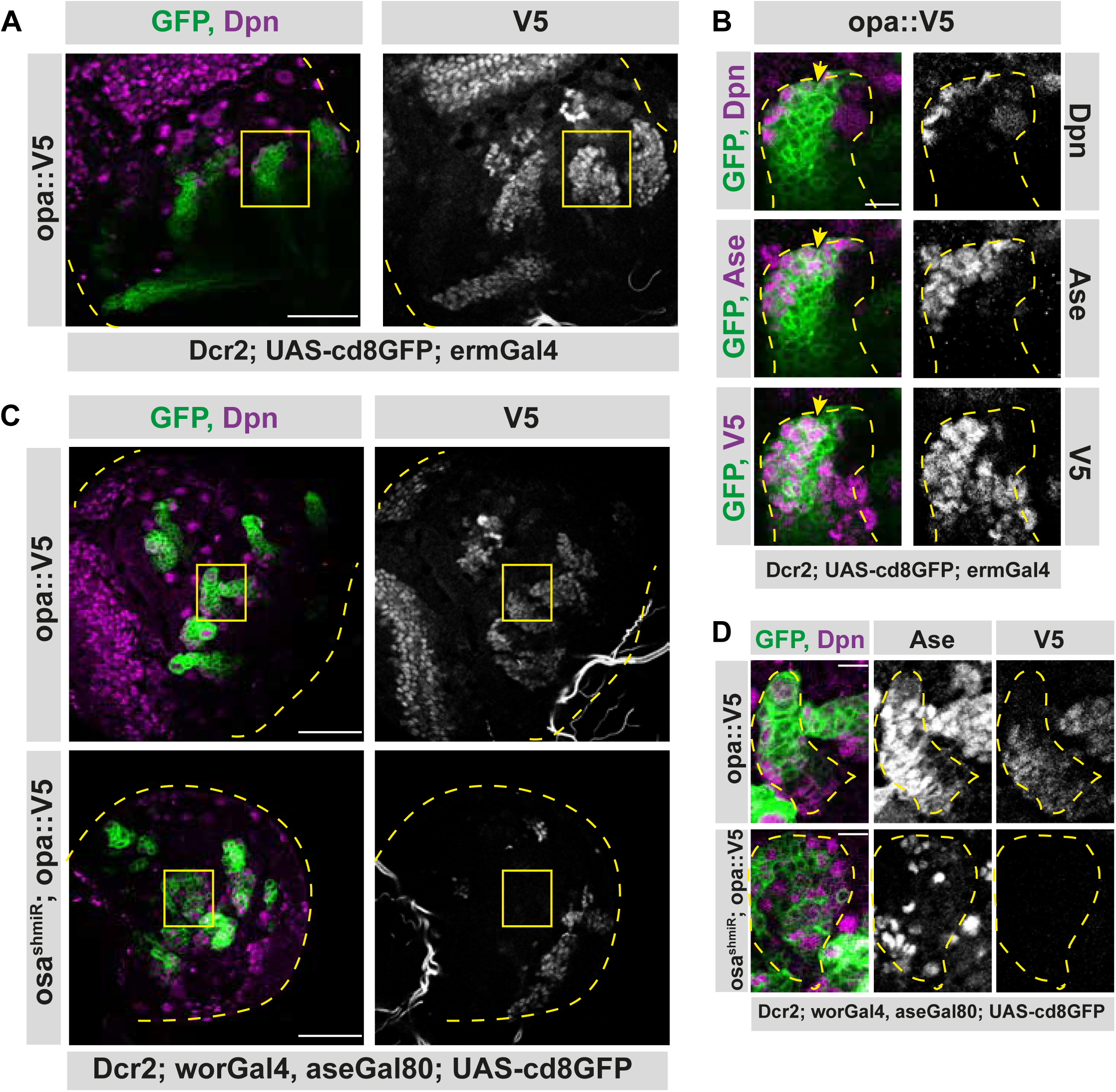
Osa initiates the expression of opa in INPs. (**A**) Opa is expressed in type II lineages. Overview image of brain lobe expressing endogenously V5-tagged opa, stained for Dpn and V5 antibodies, lobes are outlined with yellow dashed lines, scale bar 50 μm, (induced with ermGal4, marked with membrane bound GFP). (**B**) Close-up images of (**A**) marked with yellow square, stained for Dpn, Ase, and V5 antibodies. Type II lineage is outlined with yellow dashed lines, scale bar 10 μm, (induced with ermGal4, marked with membrane bound GFP). Yellow arrow is marking the start of opa expression. (**C**) Opa is induce directly by Osa. Overview images of brain lobes expressing opa-V5 alone or along with osa shmiR in type II lineages, stained for Dpn and V5 antibodies, lobes are outlined with yellow dashed lines, scale bar 50 μm, (induced with worGal4, aseGal80, marked with membrane bound GFP). Osa knock-down causes the loss of V5 expression in type II lineages. (**D**) Close-up images of (**C**) marked with yellow square, stained with Dpn, V5 and Ase antibodies. Type II lineage is outlined with yellow dashed lines, scale bar 10 μm, (induced with worGal4, aseGal80, marked with membrane bound GFP). Knock-down of Osa causes higher numbers of Dpn^+^ cells which are V5^-^.

**Figure5 supplement1.**
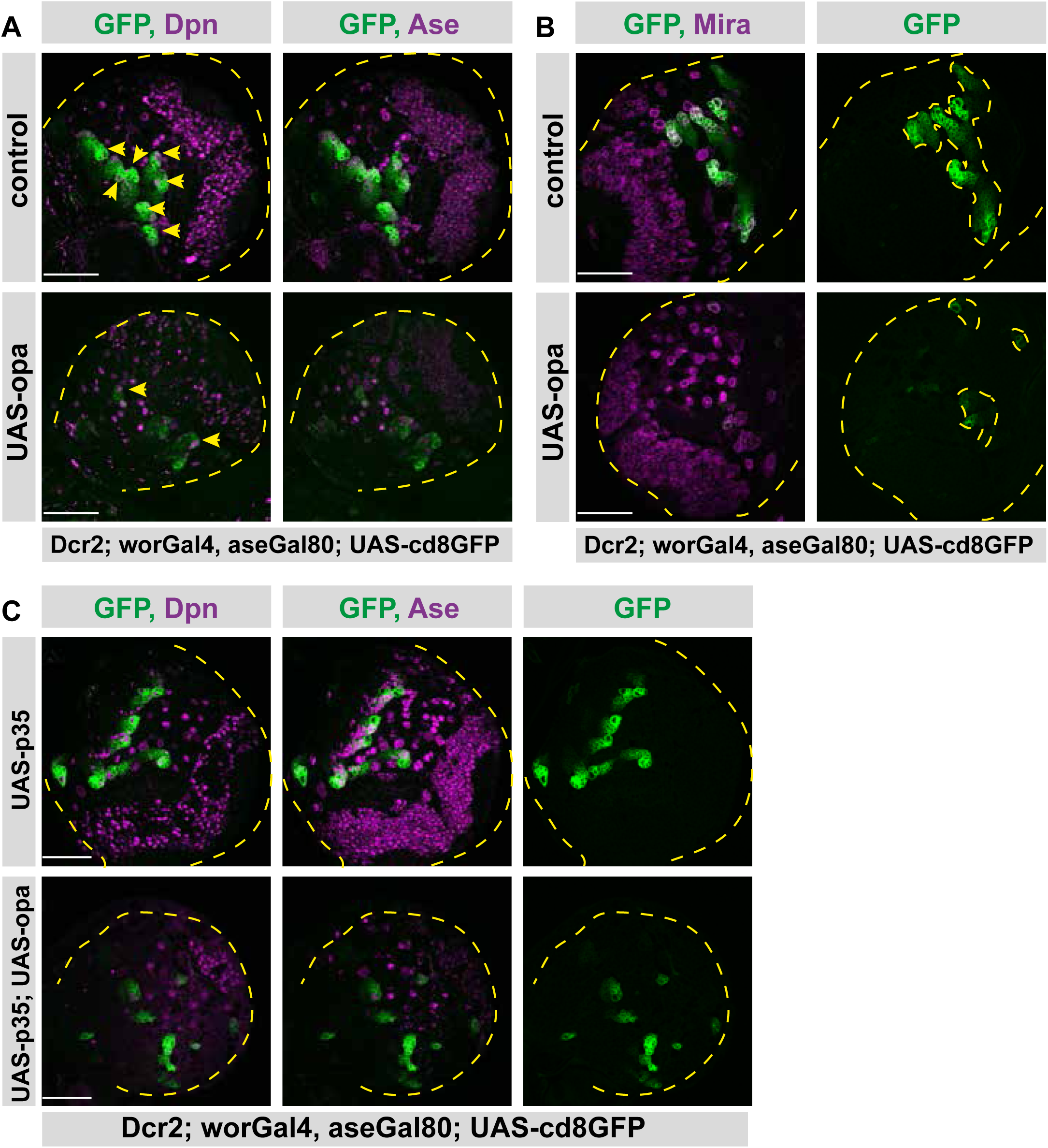
Opa overexpression causes shorter type II lineages. (**A**) Overview images of brain lobes, control or type II lineage-specific opa overexpression, stained for Dpn, and Ase antibodies, lobes are outlined with yellow dashed lines, yellow arrowheads mark Dpn positive type II NB lineages, one lineage is invisible in this z-plane, scale bar 50 μm, (induced with worGal4, aseGal80, marked with membrane bound GFP). (**B**) Overview images of brain lobes, control or type II lineage-specific opa overexpression, stained for Mira antibody, lobes and type II lineages are outlined with yellow dashed lines, scale bar 50 μm, (induced with worGal4, aseGal80, marked with membrane bound GFP). (**C**) Overexpression of apoptosis inhibitor p35 in type II lineages is not sufficient to prevent type II lineage loss upon opa overexpression. Overview images of brain lobes overexpressing opa alone and together with p35, stained for Dpn and Ase antibodies, lobes are outlined with yellow dashed lines, scale bar 50 μm, (induced with worGal4, aseGal80, marked with membrane bound GFP).

**Figure6 supplement1.**
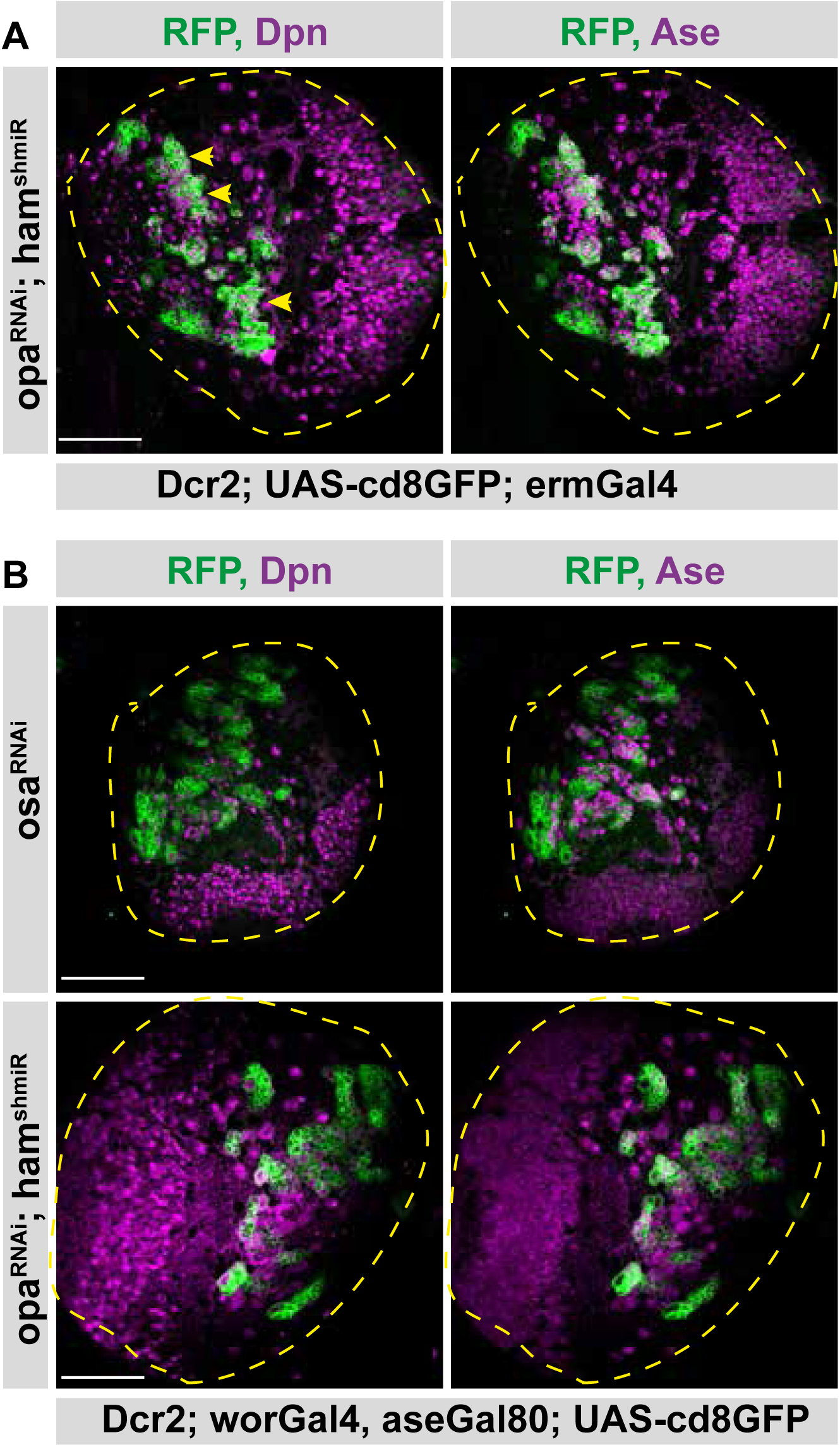
Opa and hamlet cannot recapitulate Osa knock-down phenotype. (**A**) Overview images of opa and ham RNAi expressing brains in type II lineages are stained for Dpn and Ase antibodies, brain lobes are outlined with yellow dashed lines, yellow arrowheads mark lineages with overproliferation, scale bar 50 μm, (induced with ermGal4, marked with membrane bound GFP). (**B**) Overview images of osa RNAi, and opa/ham double RNAi expressing brains in type II lineages are stained for Dpn and Ase antibodies, brain lobes are outlined with yellow dashed lines, scale bar 50 μm, (induced with worGal4, aseGal80, marked with membrane bound GFP).

